# Increased Quadriceps Intermuscular Adipose Tissue in Chronic Liver Disease is Associated with an Altered Muscle Transcriptome compared with Healthy Age Matched Controls

**DOI:** 10.1101/2025.09.29.678514

**Authors:** J I Quinlan, T Nicholson, C Bell, A Dhaliwal, F Williams, S L Allen, S Choudhary, A V Rowlands, A M Elsharkawy, M J Armstrong, C A Greig, J M Lord, Simon W Jones, L Breen

## Abstract

Muscle fat infiltration (i.e., myosteatosis) is an important index of muscle health and pathology in disease states, such as chronic liver disease (CLD). However, it is unknown whether myosteatosis differs due to anatomical location. Further, the mechanistic implications of divergent intermuscular adipose tissue (IMAT) deposition remains unknown in patients with CLD.

33 patients with CLD (55.0±10.5years) and 17 healthy controls (HC) (49.6±15.4years) were recruited. Quadriceps IMAT was estimated between 20-80% of muscle length and psoas IMAT at L3 via MRI. Body composition, muscle strength and habitual activity were assessed. Fasted vastus lateralis muscle biopsies were collected and subjected to RNA sequencing analysis. https://ClinicalTrials.gov identifier: NCT04734496.

IMAT was greater in CLD compared with HC in both quadriceps and psoas (P<0.0001). Quadriceps IMAT positively correlated with BMI (r=0.62), body fat (r=0.65), age (r=0.36) and negatively correlated with maximal knee extensor strength (r=-0.45) and habitual physical activity (r=-0.50) in CLD and HC. 169 differentially expressed genes (DEGs) were identified in low IMAT CLD vs HC, and 178 DEGs were identified in high IMAT CLD vs HC. CLD patients with high and low IMAT exhibited defined expression profiles, with only 39 DEGs (representing 12.7%) in common. Pathway analysis of DEGs revealed enrichment of atrophic and pro-inflammatory pathways in the high IMAT group.

IMAT is greater in patients with CLD compared to HC, irrespective of anatomical location and is associated with reductions in muscle function and may be exacerbated by physical inactivity. IMAT appears to alter the muscle transcriptome in COLD, potentially exacerbating muscle loss.

## Introduction

Sarcopenia, i.e., the progressive loss of muscle mass and function occurs with ageing(Cruz-Jentoft *et al*., 2019), however it may also occur secondary with existing disease, e.g., chronic kidney disease (CKD) (Sabatino *et al*., 2021), inflammatory bowel disease(Dhaliwal *et al*., 2021*a*), and chronic liver disease (CLD)(Allen *et al*., 2020). Aside from the loss of absolute muscle mass, sarcopenia is associated with reduced muscle quality, which is often defined as a reduction in strength per unit of muscle(Naimo *et al*., 2021), and as such is an important consideration in overall muscle health, as highlighted in the 2019 European Working Group on Sarcopenia in Older People consensus(Cruz-Jentoft *et al*., 2019). One such factor affecting muscle quality is the infiltration of adipose tissue into skeletal muscle, known as myosteatosis. This infiltration can be characterised as either intermuscular (IMAT) (i.e., fat deposition between muscle fascicles or beneath the muscle fascia) or intramuscular adipose tissue (i.e., fat deposition within muscle fascicles and fibres)(Correa-de-Araujo *et al*., 2020).

IMAT is known to increase with age(Buford *et al*., 2012; Akima *et al*., 2015; Hogrel *et al*., 2015; Ogawa *et al*., 2021) and can increase the secretion of local proinflammatory adipokines(Carobbio *et al*., 2011) and cytokines into the surrounding skeletal muscle(Fontana *et al*., 2007; Beasley *et al*., 2009; Delmonico *et al*., 2009; Addison *et al*., 2014), and systemically (Addison *et al*., 2014; Persson *et al*., 2022). This pro-inflammatory state is postulated to negatively impact maintenance of muscle mass, strength and function and is a likely contributor to the sarcopenia severity (Wilhelm *et al*., 2014; Zamboni *et al*., 2019; Biltz *et al*., 2020; Akazawa *et al*., 2021*b*). Further, myosteatosis has negative systemic effects including associations with diabetes and insulin resistance(Goodpaster *et al*., 2003; Boettcher *et al*., 2009), acts as an independent cardiovascular risk factor(Therkelsen *et al*., 2013) and is associated with an increased mortality risk(Reinders *et al*., 2016; Horii *et al*., 2020). Unsurprisingly, myosteatosis is heightened in the presence of chronic disease, including CLD(Kamiliou *et al*., 2024) and its clinical importance is being increasingly recognised(Henin *et al*., 2024). Previous studies have shown that excess IMAT occurs in the L3/L4 muscle groups of patients with cirrhosis(Montano-Loza *et al*., 2016; Bhanji *et al*., 2018) and the thigh muscles of patients with non-alcoholic fatty liver disease(Linge *et al*., 2021). Further, we have recently shown that quadriceps IMAT is increased in patients with end stage liver disease compared with healthy age-matched controls(Quinlan *et al*., 2023) and that this excess infiltration is associated with accelerated muscle and epigenetic ageing(Nicholson *et al*., 2024). It is therefore highly likely that myosteatosis plays a role in reducing muscle quality and sarcopenia progression in CLD patients. However, the thorough investigation of the pathophysiology of myosteatosis for skeletal muscle health is lacking, and it is unknown whether myosteatosis differs due to anatomical location in patients with CLD (i.e., different muscle groups and/or locations length). Specifically, the psoas muscle is commonly utilised to screen for reduced muscle mass and/or quality in clinical settings (REFS); however it is unknown whether this is the most suitable location to quantify myosteatosis in CLD. In line with this concept, our previous data showed greater declines in quadriceps muscle mass compared with psoas in patients with CLD(Quinlan *et al*., 2023). Therefore, an improved detailed understanding is of particular interest due to myosteatosis acting as a prognostic factor for morbidity, mortality and adverse perioperative outcomes in patients with end stage liver disease(Montano-Loza *et al*., 2016; Czigany *et al*., 2020)

This study therefore had three main aims: 1) To compare IMAT of the psoas, quadriceps muscles (i.e., rectus femoris, vastus lateralis, vastus medialis and vastus intermedius) and quadriceps location (distal to proximal) between patients with CLD and age/sex-matched healthy control participants. 2) to explore the correlation between IMAT and measures of muscle function, muscle strength and physical activity. 3) To explore the underlying muscle transcriptomic implications of IMAT in CLD.

## Methods

### Study population

33 Patients with CLD (Median age 55±10.5 years) were recruited from the liver transplant waiting list outpatient clinic at the Queen Elizabeth University Hospital Birmingham (UK) between 2019 and 2020. The CLD cohort was a component of the prospective observational study, the Evaluation of Sarcopenia in Inflammatory Disease (ESCID) (Dhaliwal *et al*., 2021*b*). In addition to the patient group, we also recruited 17 (Median age 48.9±15.6 years) healthy (i.e., no comorbidities, non-obese) age- and sex-matched control participants (HC). The study was approved by the Health Research Authority - West Midlands Solihull Research Ethics Service Committee Authority (REC reference: 18/WM/0167) and the study has been performed in accordance with the ethical standards laid down in the 1964 Declaration of Helsinki and its later amendments. All patients provided written informed consent. (https://ClinicalTrials.gov Identifier: NCT04734496).

### Intermuscular Adipose Tissue (IMAT)

IMAT was estimated via the two-point Dixon sequence method similar to previous (Ogawa *et al*., 2020; Quinlan *et al*., 2023) with offline analysis via Horos software (version 3.3.6). IMAT was calculated on the anatomical cross sectional area (ACSA) of the whole quadriceps at various anatomical locations along the full length of the muscle, i.e., 80%, 60%, 50%, 40% and 20% of muscle length with 100% being the appearance of the lesser trochanter, and 0% reflecting the appearance of the patella (Figure 1). Measures of IMAT were also completed on the ACSA of each individual quadricep muscle (i.e., vastus lateralis (VL), vastus medialis (VM), vastus intermedius (VI) and rectus femoris (RF)) at 50% of muscle length only. In addition to measures of quadriceps IMAT, psoas IMAT was also assessed at the level of L3. This analysis was only completed with n=31 for CLD and n=17 for HC due to image availability. For all muscles and locations, manual offline segmentation of the ACSA was completed on both the ‘fat only’ and ‘water only’ fractions. Mean signal intensity of the appropriate ROI was calculated for each fraction and IMAT percentage was then calculated as per equation 1. Data herein for the quadriceps are generated from the dominant leg and presented as grouped mean ± standard deviation (SD). Data generated for psoas IMAT is the average of left and right muscles and presented as grouped mean ± SD.

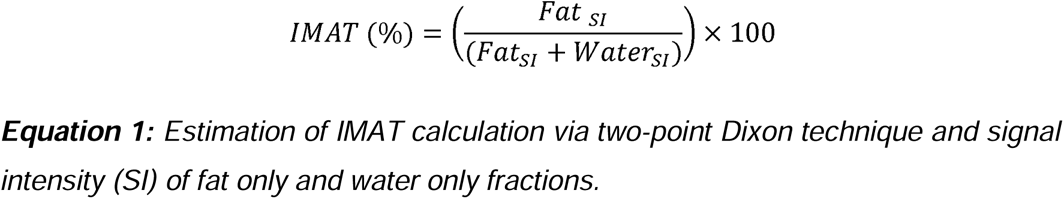

**Figure 1:**
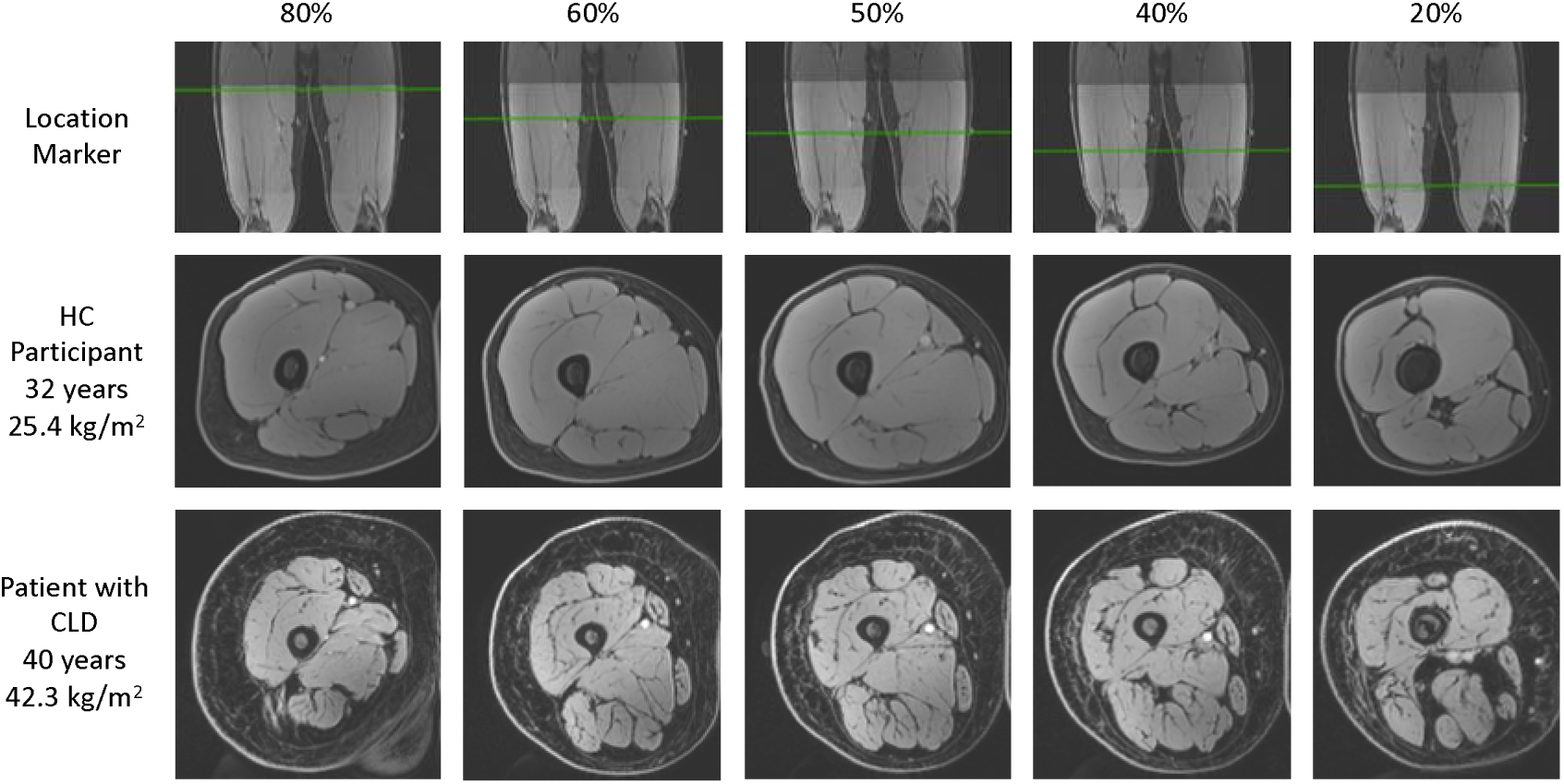
Illustrative figure demonstrating the location of each thigh scan and comparison of scans obtained from healthy control (HC) and patient with CLD.

### Isokinetic Quadriceps Strength

Measurements of unilateral peak isokinetic torque for the knee extensors (quadriceps) was conducted on the Biodex Medical System 3 (Biodex Medical Systems, New York, United States). The assessment protocol consisted of 5 consecutive maximal isokinetic leg extension contractions (60 deg•s^-1^) of the non-dominant leg. The non-dominant limb had to be utilised as muscle biopsies were collected from the dominant leg prior to this assessment as part of the ESCID study(Dhaliwal *et al*., 2021*b*). Participants with CLD underwent a familiarisation session within 2 weeks prior to the ESCID study, whereby we observed no difference in peak torque between dominant and non-dominant leg, thus validating the use of the non-dominant limb herein. Peak isokinetic torque was defined as the highest recorded torque value during the 5 completed contractions.

### Physical activity

n order to assess habitual physical activity, participants were provided with a wrist worn GENEActiv® (Activinsights, Cambridge UK) which was worn for a minimum of 3 days and maximum of 14 days prior to the visit. Extracted accelerometer files were processed and analysed with an open-source R-package, GGIR (Version 2.5-0, http://cran.r-project.org) (Rowlands *et al*., 2016; van Hees & Migueles, 2025). Data were excluded if post-calibration error was greater than 0.01 *g,* fewer than 3 valid days (≥16 hours wear) were obtained, or wear data was not present for each 15 minutes period of the 24 hour cycle. The average acceleration of movement, a proxy for total activity, was calculated for each valid day and consequently averaged across all valid days.

### Body Fat Percentage

Body composition, including body fat percentage, was estimated via bioelectrical impedance analysis (Tanita T5896). Briefly, participants stood bare foot on the device for approximately 30 seconds until instructed by the device. A readout including BMI, BMR, body fat, free fat mass, and total body water was obtained and subsequently body fat percentage was used herein for analysis.

### Muscle Biopsy

All participants reported to the laboratory in a fasted state (from 06:00 on the day of visit) and were asked to refrain from the consumption of caffeine on the morning of the trial. In addition, participants were asked to refrain from strenuous exercise for 24 h prior to their laboratory visit. A skeletal muscle biopsy was obtained from the vastus lateralis of the dominant leg using a Bergström needle (Quinlan *et al*., 2022) and immediately snap frozen in liquid nitrogen. The samples were stored at −80 °C until analysis.

### RNA isolation

Muscle tissue biopsies (∼10□mg) were first homogenized in RLT buffer (Qiagen, Manchester, UK) supplemented with Beta-mercaptoethanol (β-ME) utilizing a Qiagen Tissue Ruptor (Qiagen, Manchester, UK). RNA was then isolated and DNase-treated using a commercially available kit following the manufacturer’s protocol (RNeasy Fibrous Tissue Mini Kit, Qiagen Manchester, UK). RNA quantity and RNA integrity (all samples, RIN>7) were measured utilizing a Bioanalyzer (Agilent, CA, USA).

### Bulk RNA sequencing

For muscle tissue samples, library preparation was performed by the Genomics Facility at the University of Birmingham using a QuantSeq 3′ kit (Lexogen), with libraries sequenced on an Illumina NextSeq 500 platform. Sequencing read quality checks were performed using fastQC, and reads were trimmed using Trimmomatic. Reads were mapped to the hg38 reference human genome using Star Aligner. Differential expression analysis was determined using the DESeq2 R Bioconductor package. Pathway analysis was performed utilising Ingenuity pathway analysis (IPA; Qiagen).

### Muscle transcriptome group comparison

For the purpose of the muscle transcriptomic analysis, participants were stratified into quartiles according to the full data set IMAT percentage (i.e., HC and CLD combined), with Q1 classed as low IMAT and Q4 as high IMAT, in turn creating three groups: Healthy patients with low IMAT (HC), patients with CLD and low IMAT (IMAT Low) and patients with CLD and high IMAT (IMAT High). 4 patients with available muscle transcriptomic data were subsequently selected from each group, with groups matched for sex and age. Furthermore, HC and low IMAT groups were matched for IMAT percentage.

### Statistical analysis

All data were analysed utilising GraphPad Prism software, version 10 (La Jolla, CA, USA). Data were checked for normal distribution via the D’Agostino & Pearson test and presented as mean (standard deviation) when normally distributed and median (IQR) when non-normally distributed. Comparison between two groups were made via t-test (Mann-Whitney in the case of non-parametric). The comparison of quadriceps vs psoas IMAT, utilised the quadriceps 50% location due to it representing the mid-point of the muscle and commonly used in research. Comparisons between multiple groups or locations/muscle were assessed via two-way ANOVA and Šidák’s post hoc analysis. Correlations were assessed via Pearson r statistical test, or Spearman if non-parametric. Cohen’s d was used to calculate the effect size, where d = 0.2, 0.5 and 0.8 indicates a small, medium, and large effect, respectively. The level of significance was set at p<0.05 throughout apart from the cutoff for differentially expressed genes (DEGs) where the cut-off was defined as an adjusted p-value (q-value) of <0.05.

### Funding

This work is supported by the National Institute for Health Research Birmingham Biomedical Research Centre at the University Hospitals Birmingham NHS Foundation Trust and the University of Birmingham (BRC-1215-20009). AR is supported by the National Institute for Health Research (NIHR) Leicester Biomedical Research Centre and NIHR Applied Research Collaborations East Midlands. The views expressed are those of the authors and not necessarily those of the NHS, the NIHR or the Department of Health and Social Care.

## Results

### Demographics

Demographics were collated for all participants, with no difference in age or height between groups. However, whilst body weight (91.4kg (23.1) vs 73.3kg (12.6), P<0.01) and BMI (30.2kg/m2 (6.9) vs 24.9kg/m2 (3.3), P<0.01) was significantly higher in patients with CLD, body fat percentage was similar between groups (30.8% (9.9) vs 27.3 % (7.9) for CLD and HC respectively). For transcriptome analysis, groups were counterbalanced for age (54.5, 58 and 55 years for CLD low IMAT, CLD high IMAT and HC, respectively) and IMAT percentage where appropriate (7.8% vs 6.9% for CLD low IMAT and HC, respectively).

### The effect of anatomical location on measures of IMAT

Psoas IMAT at the level of L3 was significantly greater than quadriceps IMAT measured at 50% of muscle length (14.8 (5.7) vs 10.1 (4.0), P<0.0001, d=0.94) when CLD and HC data were combined (i.e., psoas vs quad). Regarding the quadriceps alone, two-way ANOVA revealed a main effect of anatomical location (i.e., distal to proximal) (P<0.0001) and condition (i.e., CLD vs HC) (p<0.0001) for whole quadriceps IMAT, in addition to a significant interaction (i.e., location x condition) (P<0.01) (Figure 2A). Šidák’s post hoc comparison revealed that quadriceps IMAT was greater in CLD at all anatomical locations (20-80% of quadricep muscle length) compared to HC (Supplementary table 1). Further, in both CLD and HC, quadriceps IMAT was greatest at 20% of muscle length (Supplementary table 2).

**Figure 2:**
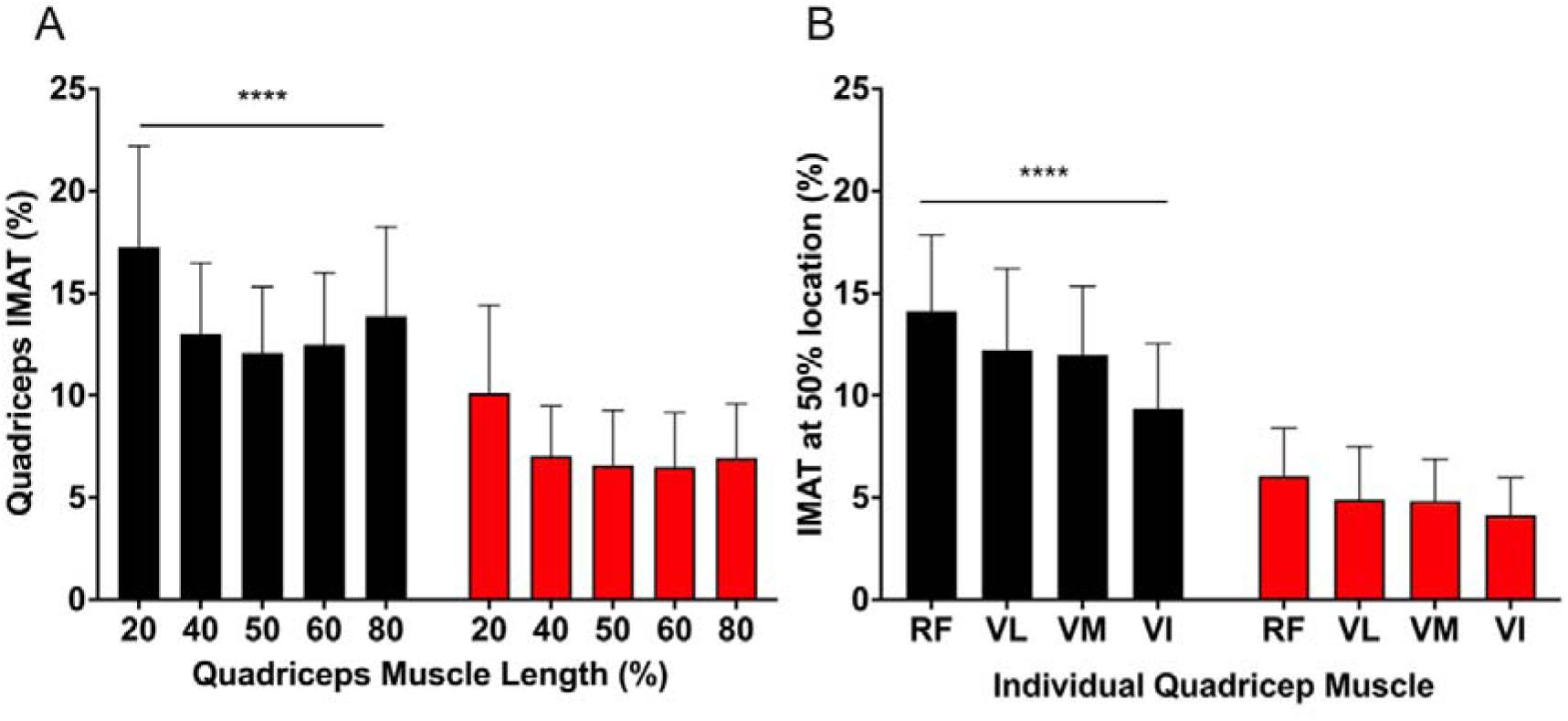
Comparison of quadriceps IMAT across the length of the muscle (A) and between individual muscles at the 50% location (B) in patients with CLD (black) and healthy control participants (red). Significance defined by **** indicates an effect of condition (i.e., CLD vs HC), P<0.0001.

### The effect of individual quadriceps muscle

When comparing individual quadricep muscles 50% of quadriceps length (i.e., VL, VI, VM, RF), a two-way ANOVA revealed a main effect for muscle (P<0.0001) and condition (P<0.0001) with a significant interaction (i.e., muscle x condition) (P<0.01) (Figure 2B). Šidák’s multiple comparisons revealed IMAT was significantly greater in the muscle of patients with CLD compared with HC (Supplementary table 3). Further, differences between individual muscles were seen in CLD but not in HC (Figure 2B), specifically in CLD RF IMAT was greater than VL (P<0.01), VM (P<0.001) and VI (P<0.0001), and both VL (P<0.0001) and VM (P<0.0001) greater than VI (Supplementary table 3).

### Correlations between IMAT and measures of body composition and physical function

Pearson r correlation showed significant positive correlations between quadriceps IMAT (at 50% muscle length) and BMI (r=0.62, P<0.0001, n=50), body fat percentage (r=0.65, P<0.0001, n=50) and age (r=0.36, P<0.01, n=50) (Figure 3A, C, E). However, psoas IMAT did not correlate with BMI, body fat percentage or age (P>0.05 for all) (Figure 3B, D, F). Negative correlations were seen between quadriceps IMAT and both maximal knee extensor strength (r=-0.45, P<0.01, n=50) (Figure 4A) and habitual physical activity (r=-0.50, P<0.001, n=42) (Figure 4B). Similar correlations were seen between psoas IMAT and maximal knee extensor strength (r=-0.45, P<0.001, n=48) (Figure 4C) and habitual physical activity (r=-0.51, P<0.01, n=41) (Figure 4D).

**Figure 3:**
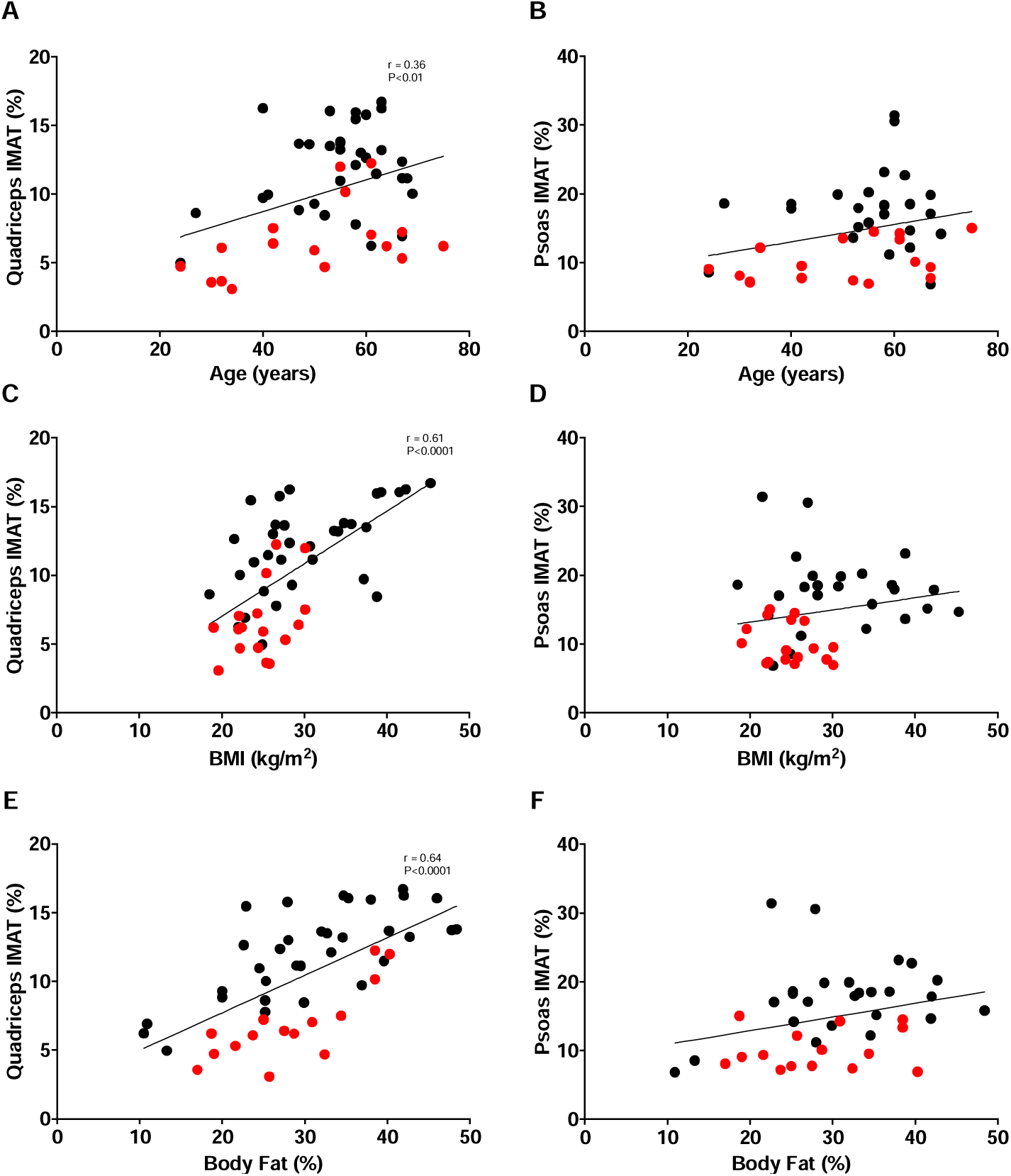
Correlation between quadriceps IMAT or psoas IMAT and age (A,B), BMI (C,D) and body fat percentage (E,F) in patients with CLD (black) and healthy control participants (red). Significant correlations indicated with r and P value, whereas absence indicates no significant correlation.

**Figure 4:**
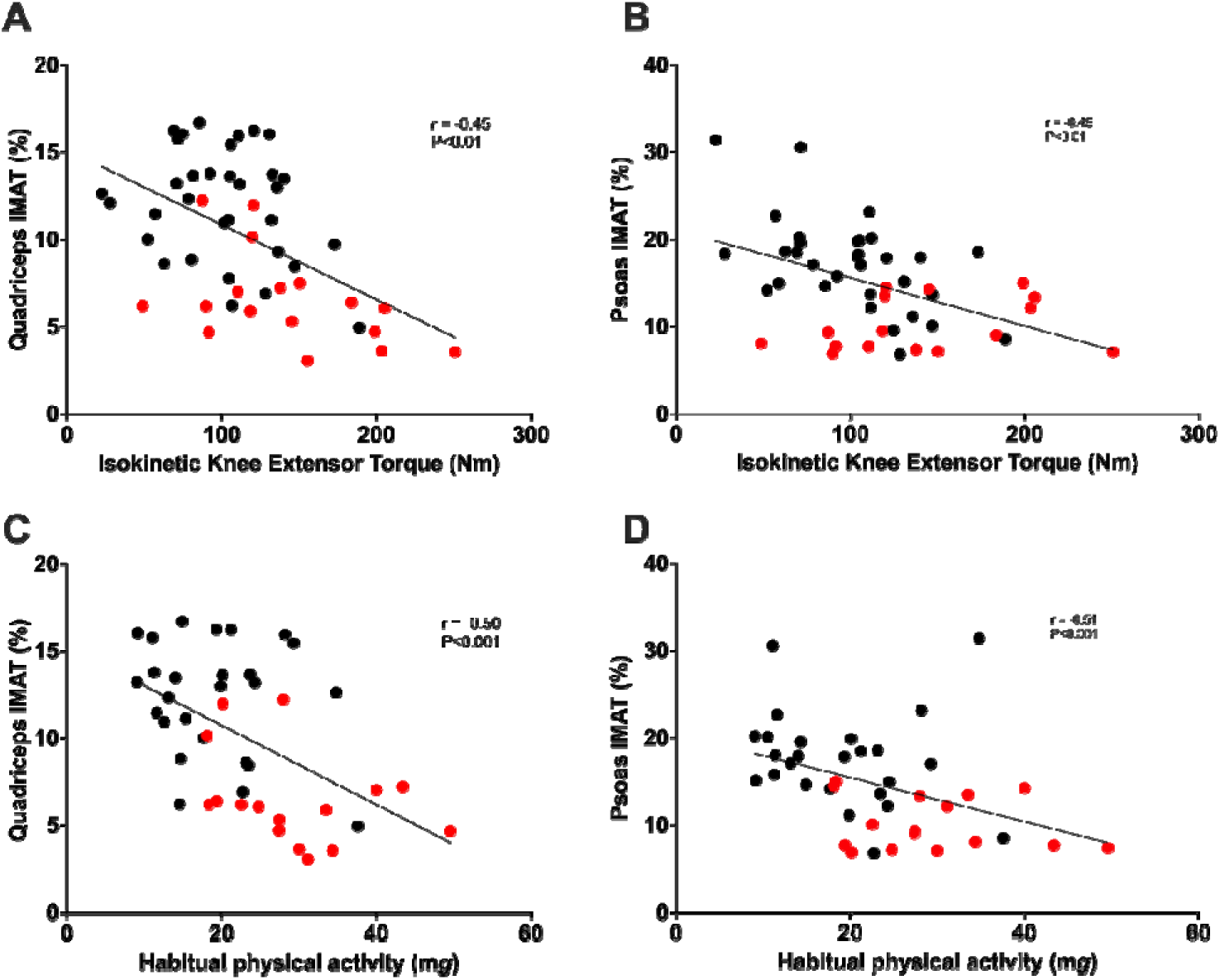
Correlations between quadriceps IMAT or psoas IMAT and maximal isokinetic knee extensor torque (A,B), and habitual physical activity (C,D) in patients with CLD (black) and healthy control participants (red). Significant correlations indicated with r and P value, whereas absence indicates no significant correlation.

### IMAT is associated with an altered muscle transcriptome in CLD patients that differs with increasing IMAT percentage

Having identified the presence of significantly greater IMAT in the muscle of CLD patients, we next sought to determine the potential impact of IMAT in CLD patient on muscle.RNAseq identified 169 and 178 differentially expressed genes (DEGs, q-value <0.05) in the skeletal muscle of CLD patients with low IMAT or high IMAT vs HCs respectively. Low and high IMAT individuals exhibited defined expression profiles, with only 39 DEGs (12.7%) common between groups. Pathway analysis of DEGs revealed greater enrichment of pathways associated with atrophy (degradation of branched chain amino acids) and inflammation (neutrophil degranulation) in the high IMAT group and altered metabolism in the low IMAT group (AMPK signalling, Mitochondrial translation and Glutathione synthesis).

**Figure 5:**
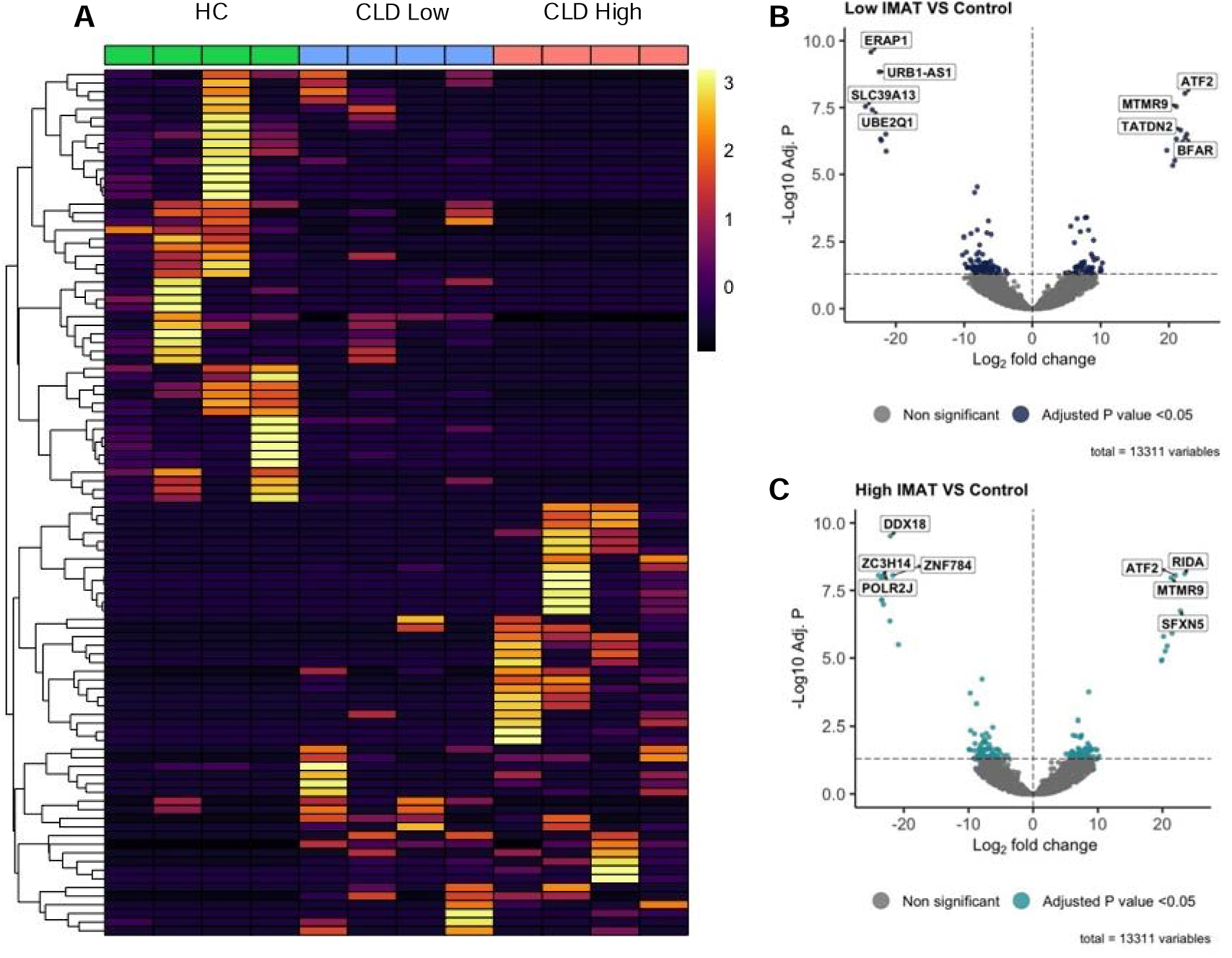
Heatmap displaying differentially expressed in patients with CLD and low (CLD Low) and high IMAT (CLD High) in comparison with healthy controls (HC) (A). Volcano plot showing significantly different genes between HC and patients with CLD and low IMAT (B) or patients with CLD and high IMAT (C)

**Figure 6:**
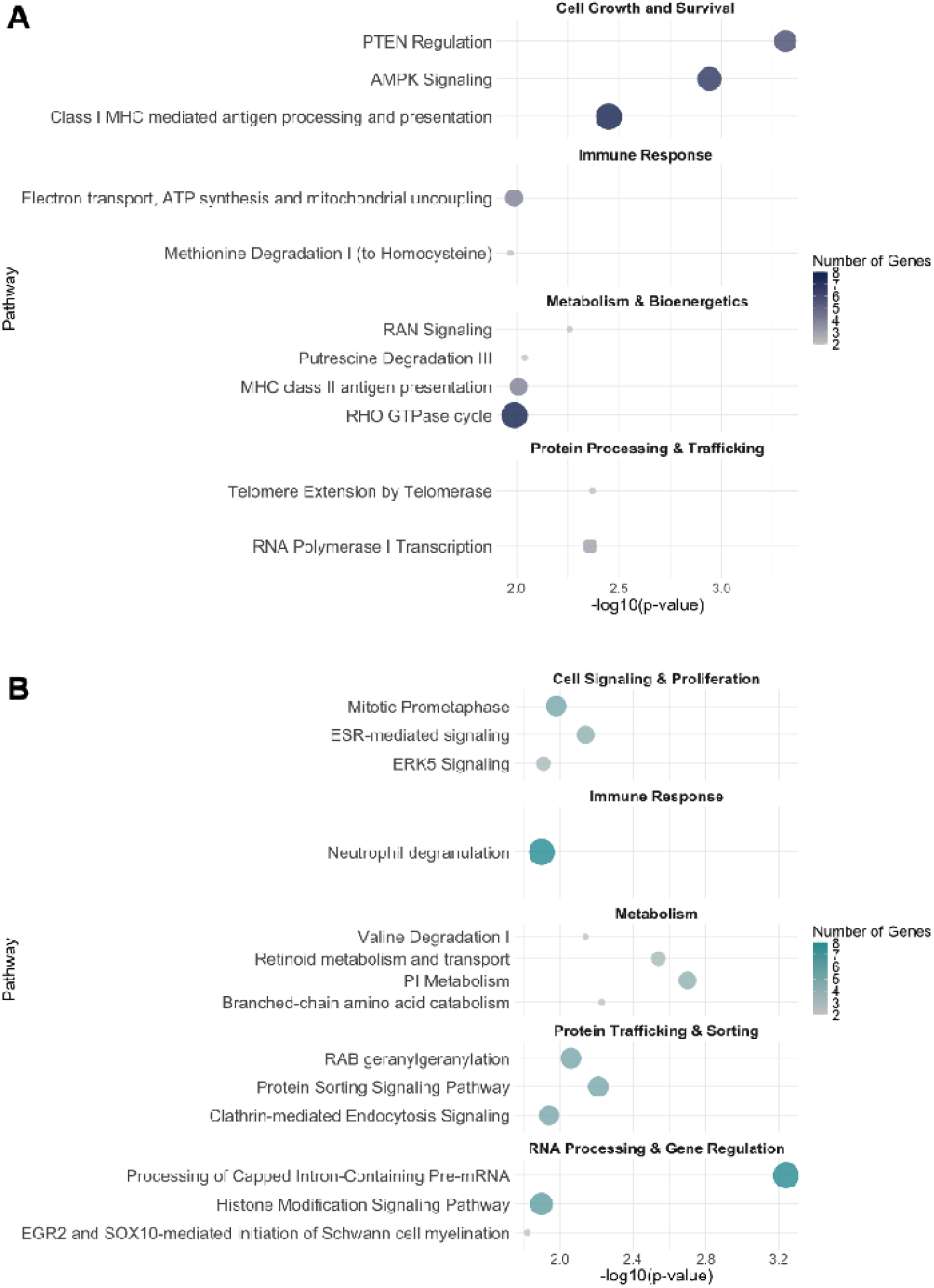
Pathway analysis demonstrating the top differentially expressed pathways in patients with CLD and low (A) and high IMAT (B) in comparison with health controls (HC).

## Discussion

Herein we confirm that IMAT is greater in CLD patients compared to healthy age and sex matched controls; irrespective of anatomical location. The presence of myosteatosis in CLD appeared to be associated with an altered muscle transcriptome compared with HC, with pathways concomitant with muscle metabolism, amino acid degradation and inflammation all negatively impacted.

The first aim of this study was to investigate whether IMAT values of muscles from different anatomical locations differed to one another within a CLD cohort, specifically, muscles at the level of L3 and the quadriceps due to their utility in both the clinical and scientific research fields. Following analysis of the full dataset (i.e., HC and CLD combined), we observed that psoas IMAT at the level of L3 was significantly greater compared to quadriceps IMAT measured at 50% of muscle length, demonstrating that levels of IMAT may differ due to their anatomical location. This finding is of particular relevance for this patient group given the association between psoas IMAT and the increased risk of mortality (HR of 2.94) following hepatocellular carcinoma surgery (Hamaguchi *et al*., 2015). Nonetheless, when comparing IMAT values between CLD and HC (i.e., HC psoas vs CLD psoas and HC quadriceps vs CLD quadriceps), greater differences in IMAT were seen between the quadriceps compared with the psoas (81% vs 69% respectively). This finding suggests that the quadriceps may be more susceptible to myosteatosis than truncal muscles in patients with CLD. This is line with our previous research where we highlighted that muscle loss was greater in the lower limbs compared to truncal muscles in patients with CLD(Quinlan *et al*., 2023). Whilst the combination of quadriceps muscle loss and increased IMAT will have an inherent negative impact on muscle function; it has been suggested that the negative impact of IMAT is greater, with increased quadriceps IMAT associated with poorer recovery of activities of daily living following hospital admission than reduced muscle mass(Akazawa *et al*., 2021*a*).

IMAT impacts muscle function and strength in older individuals(Wilhelm *et al*., 2014; Akazawa *et al*., 2017, 2020) and it would be logical to suggest the same relationships would be observed in CLD patients. Indeed, both quadriceps and psoas IMAT were negatively correlated with maximal knee extensor torque. Whilst this is an expected result for the quadriceps, it is surprising this correlation was also observed with the psoas, given the lack of direct muscular involvement of the psoas in this assessment. However, in consideration of the negative correlation between both quadriceps and psoas IMAT and habitual physical activity; it is possible that psoas IMAT is more reflective of wider health status rather than a specific indication of lower body function. To this end, we postulate that the reduction in physical activity as a consequence of CLD(Quinlan *et al*., 2023) is in part responsible for the increase in IMAT seen within this patient group. Collectively, this highlights the clinical need to proactively address reductions in muscle quality, as well as muscle mass in patients with CLD.

The final aim of this study was to understand the mechanistic implications of IMAT through comparison of the muscle transcriptome of healthy individuals and CLD patients with either high or low IMAT. Our data revealed differences between the patient and non-patient groups (178 and 169 DEGs for low and high IMAT respectively), with the CLD groups demonstrating distinct expression profiles with only 12.7% common genes between groups (39 DEGs). Compared with HC, patients with CLD and high quadriceps IMAT, exhibited altered expression of genes related to amino acid metabolism (e.g., branched-chain amino acid catabolism and valine catabolism) and inflammation (neutrophil degranulation). When combined with altered ESR-mediated signalling and oxidative stress, these factors may evoke a catabolic environment within the muscle, in turn propagating the accelerated muscle loss seen in CLD (Quinlan *et al*., 2023).

Whilst we may have expected differences in the muscle transcriptome between HC and high IMAT, we also observed 178 DEGs between HC and CLD patients with Low IMAT, even when matched for IMAT percentage, age and sex. This could indicate a disease-related switch of IMAT in patients with CLD, which drives pathological crosstalk with muscle. Pathway analysis revealed these DEGs to be associated with altered metabolism, i.e., AMPK signalling, mitochondrial translation, glutathione synthesis and electron transport and ATP synthesis. We have previously shown that mitochondrial respiration, coupling efficiency and mitophagy are all reduced in end stage liver disease (Allen *et al*., 2022). Importantly this reduction in energy availability would not only have implications for muscle contraction, but also for energy intense procedures such as muscle protein synthesis. These data reinforce our previous work where we found a strong correlation between IMAT in CLD and accerlerated biological ageing, even in individuals with lower BMI (i.e., <25kg/m^2^) (Nicholson *et al*., 2024). Collectively, these data build upon the notion that skeletal muscle of patients with CLD is significantly compromised through an altered expression of a multitude of transcriptomic pathways which may be exacerbated by the presence of increased IMAT. Indeed, research from our group has shown that adipose tissue can have direct implications for skeletal muscle (Nicholson *et al*., 2018; O’Leary *et al*., 2018); nonetheless, future work should aim to identify the specific mechanisms in which this adipose-muscle crosstalk occurs in patients with CLD to alter the muscle transcriptome e.g., cytokines, extracellular vesicles, increased inflammatory senescent cells.

## Conclusion

We hererin demonstrate that patients with CLD have increased IMAT, a conclusion which is independent of the anatomical location. This increased IMAT and resulting reduced muscle quality is associated with reduced knee extensor strength, which may be exacerbated by reduced physical activity. Possibly quadriceps may be impacted by IMAT to a greater extent than psoas, and more strongly correlated to age, BMI and body fat percentage. Further, this increased quadriceps IMAT in CLD is associated with an altered muscle transcriptome compared with HC, even when controlled for IMAT percentage; potentially indicating the existence of a disease driven switch in the IMAT of patients with CLD. Collectively, our data suggest a greater need to consider muscle quality in interventions aimed at improving muscle health and function in patients with CLD. Future endeavours should aim to understand the specific pathways and mechanisms responsible for eliciting this altered muscle transcriptome.

---

**Supplementary table 1:**
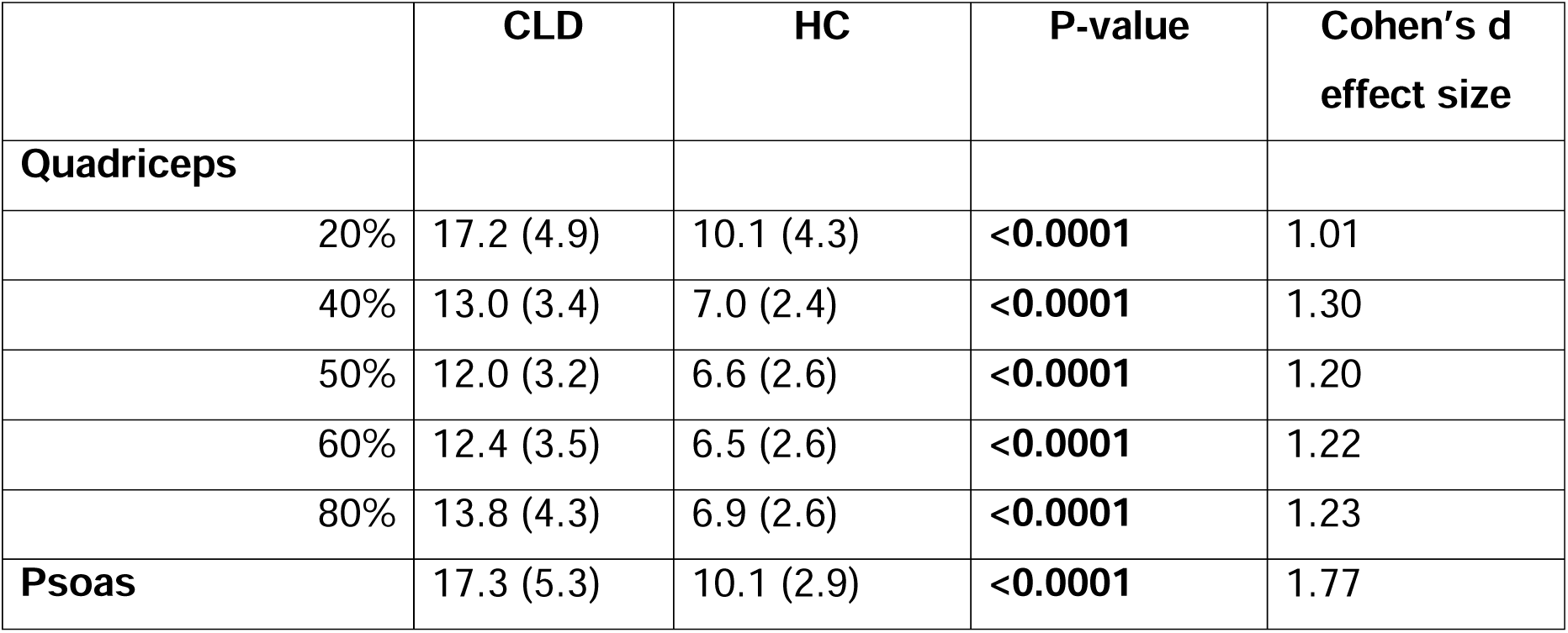
IMAT values for the quadriceps and psoas of patients with CLD (CLD) and health control participants (HC) across a range of anatomical locations. For the quadriceps, 20% reflects the most distal point and 80% most proximal point.

|  | **CLD** | **HC** | **P-value** | **Cohen's d**<br>**effect size** |
| --- | --- | --- | --- | --- |
| **Quadriceps** |  |  |  |  |
| 20% | 17.2 (4.9) | 10.1 (4.3) | **<0.0001** | 1.01 |
| 40% | 13.0 (3.4) | 7.0 (2.4) | **<0.0001** | 1.30 |
| 50% | 12.0 (3.2) | 6.6 (2.6) | **<0.0001** | 1.20 |
| 60% | 12.4 (3.5) | 6.5 (2.6) | **<0.0001** | 1.22 |
| 80% | 13.8 (4.3) | 6.9 (2.6) | **<0.0001** | 1.23 |
| **Psoas** | 17.3 (5.3) | 10.1 (2.9) | **<0.0001** | 1.77 |

**Supplementary table 2:**
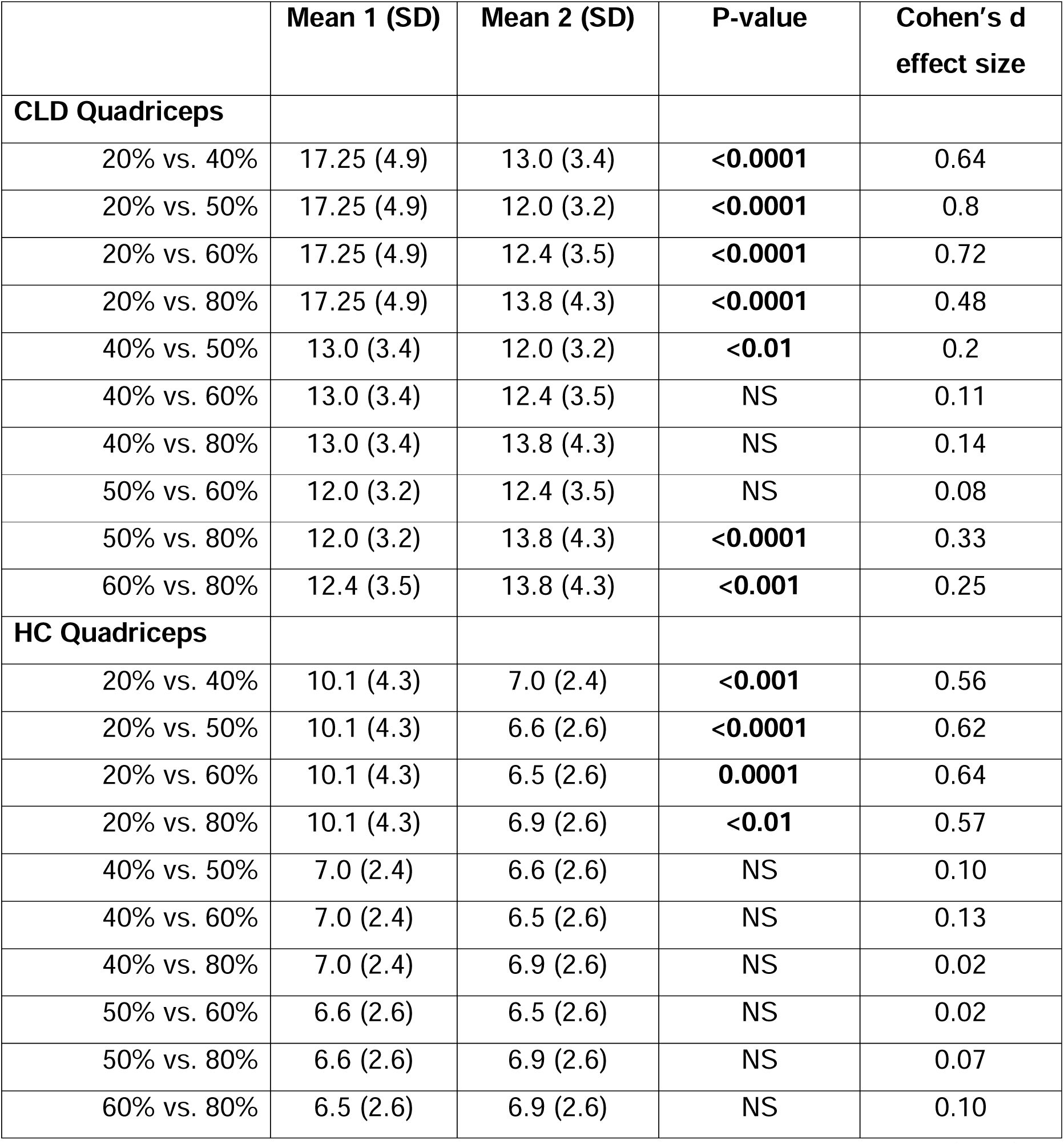
IMAT values for the quadriceps of patients with CLD (CLD) and health control participants (HC) across a range of anatomical locations with 20% being most distal and 80% most proximal.

|  | Mean 1 (SD) | Mean 2 (SD) | P-value | Cohen's d effect size |
| --- | --- | --- | --- | --- |
| **CLD Quadriceps** |  |  |  |  |
| 20% vs. 40% | 17.25 (4.9) | 13.0 (3.4) | **<0.0001** | 0.64 |
| 20% vs. 50% | 17.25 (4.9) | 12.0 (3.2) | **<0.0001** | 0.8 |
| 20% vs. 60% | 17.25 (4.9) | 12.4 (3.5) | **<0.0001** | 0.72 |
| 20% vs. 80% | 17.25 (4.9) | 13.8 (4.3) | **<0.0001** | 0.48 |
| 40% vs. 50% | 13.0 (3.4) | 12.0 (3.2) | **<0.01** | 0.2 |
| 40% vs. 60% | 13.0 (3.4) | 12.4 (3.5) | NS | 0.11 |
| 40% vs. 80% | 13.0 (3.4) | 13.8 (4.3) | NS | 0.14 |
| 50% vs. 60% | 12.0 (3.2) | 12.4 (3.5) | NS | 0.08 |
| 50% vs. 80% | 12.0 (3.2) | 13.8 (4.3) | **<0.0001** | 0.33 |
| 60% vs. 80% | 12.4 (3.5) | 13.8 (4.3) | **<0.001** | 0.25 |
| **HC Quadriceps** |  |  |  |  |
| 20% vs. 40% | 10.1 (4.3) | 7.0 (2.4) | **<0.001** | 0.56 |
| 20% vs. 50% | 10.1 (4.3) | 6.6 (2.6) | **<0.0001** | 0.62 |
| 20% vs. 60% | 10.1 (4.3) | 6.5 (2.6) | **0.0001** | 0.64 |
| 20% vs. 80% | 10.1 (4.3) | 6.9 (2.6) | **<0.01** | 0.57 |
| 40% vs. 50% | 7.0 (2.4) | 6.6 (2.6) | NS | 0.10 |
| 40% vs. 60% | 7.0 (2.4) | 6.5 (2.6) | NS | 0.13 |
| 40% vs. 80% | 7.0 (2.4) | 6.9 (2.6) | NS | 0.02 |
| 50% vs. 60% | 6.6 (2.6) | 6.5 (2.6) | NS | 0.02 |
| 50% vs. 80% | 6.6 (2.6) | 6.9 (2.6) | NS | 0.07 |
| 60% vs. 80% | 6.5 (2.6) | 6.9 (2.6) | NS | 0.10 |

**Supplementary table 3:** IMAT values for the Rectus Femoris (RF), Vastus Lateralis (VL), Vastus Medialis (VM) and Vastus Intermedius (VI) of patients with CLD (CLD) and health control participants (HC).

|  | CLD | HC | P-value | Cohen's d effect size |
| --- | --- | --- | --- | --- |
| **Quadriceps** |  |  |  |  |
| **RF** | 14.1 (3.7) | 6.0 (2.3) | **<0.0001** | 1.67 |
| **VL** | 12.2 (4.0) | 4.9 (2.6) | **<0.0001** | 1.37 |
| **VM** | 11.9 (3.3) | 4.8 (2.0) | **<0.0001** | 1.65 |
| **VI** | 9.3 (3.1) | 4.1 (1.8) | **<0.0001** | 1.30 |
| **CLD** | **Mean 1 (SD)** | **Mean 2 (SD)** |  |  |
| RF vs. VL | 14.1 (3.7) | 12.2 (4.0) | **<0.01** | 0.33 |
| RF vs. VM | 14.1 (3.7) | 11.9 (3.3) | **<0.001** | 0.40 |
| RF vs. VI | 14.1 (3.7) | 9.3 (3.1) | **<0.0001** | 0.90 |
| VL vs. VM | 12.2 (4.0) | 11.9 (3.3) | NS | 0.04 |
| VL vs. VI | 12.2 (4.0) | 9.3 (3.1) | **<0.0001** | 0.51 |
| VM vs. VI | 11.9 (3.3) | 9.3 (3.1) | **<0.0001** | 0.53 |
| **HC** |  |  |  |  |
| RF vs. VL | 6.0 (2.3) | 4.9 (2.6) | NS | 0.31 |
| RF vs. VM | 6.0 (2.3) | 4.8 (2.0) | NS | 0.35 |
| RF vs. VI | 6.0 (2.3) | 4.1 (1.8) | NS (P=0.06) | 0.58 |
| VL vs. VM | 4.9 (2.6) | 4.8 (2.0) | NS | 0.02 |
| VL vs. VI | 4.9 (2.6) | 4.1 (1.8) | NS | 0.21 |
| VM vs. VI | 4.8 (2.0) | 4.1 (1.8) | NS | 0.24 |

## Notes

### Competing Interest Statement

The authors have declared no competing interest.

